# Identification of Three Medically Important Mosquito Species Using Raman Spectroscopy

**DOI:** 10.1101/2022.05.17.492344

**Authors:** Dickson L Omucheni, Kenneth A Kaduki, Wolfgang R Mukabana

## Abstract

Accurate identification of disease vector insects is crucial when collecting epidemiological data. Traditionally, mosquitoes that transmit diseases like malaria, yellow fever, chikungunya, and dengue fever have been identified by looking at their external morphological features at different life cycle stages. This process is tedious and labour intensive.

In this paper, the potential of Raman spectroscopy in combination with Linear and Quadratic Discriminant Analysis to classify three mosquito species, namely: *Aedes aegypti, Anopheles gambiae* and *Culex quinquefasciatus*, was explored. The classification was based on the mosquitoes’ cuticular melanin. The three mosquito species represented two subfamilies of medically important mosquitoes, i.e. the Anophelinae and the Culicinae. The housefly (*Musca domestica*) was included as a ‘control’ group to assess the discrimination ability of the classifiers. This study is the first to use Raman spectroscopy to classify mosquitoes. Fresh mosquitoes were anaesthetized with chloroform, and a dispersive Raman microscope was used to capture spectra from their legs. Broad melanin peaks centred around 1400 cm^-1^, 1590 cm^-1^, and 2060 cm^-1^ dominated the spectra. Variance Threshold (VT) and Principal Component Analysis (PCA) were used for feature selection and feature extraction respectively from the preprocessed data. The extracted features were then used to train and test Linear Discriminant Analysis (LDA) and Quadratic Discriminant Analysis (QDA) classifiers.

The VT/PCA/QDA classification model performed better than VT/PCA/LDA. VT/PCA/QDA achieved an overall accuracy of 94%, sensitivity of 87% and specificity of 96%, whereas VT/PCA/LDA attained an accuracy of 85%, a sensitivity of 69% and a specificity of 90%. The success of these relatively simple classification models on Raman spectroscopy data lays the groundwork for future development of models for discriminating morphologically indistinguishable insect species.

## INTRODUCTION

Mosquitoes transmit many diseases to man, including malaria, yellow fever, chikungunya and dengue fever. Female mosquitoes are obligate blood-feeders and are responsible for transmitting these diseases (1,2). Disease transmission occurs when susceptible female mosquitoes become infected through blood-feeding, support pathogen development to maturity, and obtain the next blood meal from a susceptible individual (3). Blood meals are essential nourishment that female mosquitoes use to develop their eggs (4). Infected mosquitoes introduce disease-causing pathogens into their blood meal hosts through saliva, which is injected alongside an anticoagulant enzyme known as salivary apyrase (5).

Identification of disease vectors results in essential data that epidemiologists can use to develop strategies for disease control. Mosquito identification has traditionally been realized by observing morphological features at different life cycle stages. Identification is achieved using taxonomic keys in which individual mosquitoes are classified based on contrasting morphological features. Adults of Anopheline mosquitoes are readily separated from Culicines by their stature in resting positions. The Anophelines are known to rest with their bodies at an angle to the resting surface in a ‘head down bottom up’ posture. On the other hand, the culicines rest with their abdomens almost parallel to the resting surface. In the female adults, which are of medical importance, examination of the heads is relied on in distinguishing Anophelines from Culicines. The Anophelines have palps that are as long as the proboscis, usually lying close along with it. The palps in Culicines are shorter than the proboscis. Other features in *Anopheles* include the presence of a single spermatheca and dark scales on the wing veins arranged in ‘blocks’. In contrast, the Culicines have two or three spermathecae, and the dark scales on the wing veins are continuous and not arranged in distinctive areas (blocks). The genera *Culex* and *Aedes*, which include the most medically important species, are also identified by taxonomic keys. *Culex* species are recognized by their lack of ornamentation, which is conspicuous among *Aedes*, which have patterns of black and white or silvery scales on the thorax, abdomen and legs. In addition, the tip of the *Culex* abdomen is not pointed as it is in *Aedes* species (6).

The use of taxonomic keys by observation is generally a tedious and labour-intensive process, hence the need to develop advanced tools. The most advanced and recent techniques for identifying mosquitoes are molecular techniques. These tools focus on identifying morphologically indistinguishable species such as those within the *Anopheles gambiae* complex. Molecular techniques rely on the analysis of DNA (7–15). Currently, DNA analysis is achieved through Polymerase Chain Reaction (PCR) amplification (16). PCR is a technique used to select specific portions of an organism’s genome (DNA sequences) to be replicated several times to a reasonable quantity for analysis. PCR-based methods have high accuracy, specificity and sensitivity. However, PCR is time-consuming, labour-intensive, and expensive. It also requires special laboratory conditions and highly skilled personnel.

Matrix-Assisted Laser Desorption/Ionization Time of Flight Mass Spectrometry (MALDI-TOF-MS) has also been tested as an alternative method of discriminating the sibling species within the *Anopheles gambiae* complex. MALDI-TOF-MS is a technique that uses laser energy and an absorbing matrix to ionize large molecules, such as proteins, while minimizing fragmentation. Using MALDI-TOF-MS, mosquito leg protein extracts were adequate to identify mosquitoes to species level (17–19).

Spectroscopic techniques such as Near Infra-Red (NIR) spectroscopy have previously been investigated in insect taxonomic studies. These include the identification of species of beetles (20), *Drosophila* species (21) and the identification of cryptic *Tetramorium* ant species (22). The success of NIR spectroscopy in differentiating insect species has seen it deployed to the *Anopheles gambiae* complex identification problem (22,23). NIR spectroscopy probes vibrational states of molecules and provides a spectral fingerprint of the chemical compound under investigation. Cuticular lipids and hydrocarbons have been considered the main molecules that provide essential classification information in NIR spectroscopy insect taxonomic studies (24–26).

The MALDI-TOF-MS and NIR spectroscopy techniques have the advantage of being rapid compared to PCR methodologies. However, MALDI-TOF-MS involves a relatively time-consuming sample preparation process since the compound to be investigated must be extracted and embedded in a laser absorbing matrix for analysis. On the other hand, the NIR spectra are complicated by water absorption signatures. NIR spectra of fresh biological samples, which contain water molecules, are therefore challenging to interpret.

Raman spectra contain complementary molecular vibration information to spectra in the mid-IR range. The Raman technique has several advantages over NIR, including minimal sample preparation and the absence of water interference (27). It is usually integrated into microscopy for high spatial resolution and 3D mapping and can be miniaturized into portable hand-held devices. These attributes make Raman spectroscopy potentially attractive for public health applications in entomology.

Although there have been some studies that have used Raman spectroscopy in entomology, these are not widespread. These studies include analyses of the structure of honey bee wings (28), melanin in spiders (29), and bumble bees (30). Recently, Wang *et al*. (31) reported a study on mosquito age-grading that employed surface-enhanced Raman spectroscopy (SERS). To the best of our knowledge, Raman spectroscopy has not been used for mosquito taxonomy. In this paper, we demonstrate, for the first time, the capability of Raman spectroscopy in combination with machine learning tools to classify three species of mosquitoes: *Aedes aegypti, Anopheles gambiae* and *Culex quinquefasciatus*, based on their cuticular melanin signatures. We chose them to represent two subfamilies of medically important mosquitoes, i.e. the Anophelinae (Anopheline mosquitoes) and the Culicinae (Culicine mosquitoes). We explore the potential of two machine learning tools: Linear and Quadratic Discriminant Analysis, in discriminating the three groups of mosquitoes. We include the housefly (*Musca domestica*) as a ‘control’ group to test the discrimination ability of the classifiers.

## MATERIALS AND METHODS

### Mosquito Rearing and Sample Preparation

*Aedes aegypti, Anopheles gambiae* and *Culex quinquefasciatus* were reared in insectaries of the Department of Biology at the University of Nairobi. Adult mosquitoes were held in 30 × 30 × 30 cm cages in separate rooms. In each cage, the mosquitoes laid eggs in an ovicup containing a cone of filter paper placed on water. The eggs were transferred into trays filled with water, where they hatched into larvae. The larvae were fed on TetraMin^®^ baby fish food. The adults were fed on a 6% glucose solution soaked in filter paper wicks. The rooms were kept at a temperature of 27°C-28°C and 32°C-34°C for *Aedes/Culex* and *Anopheles* mosquitoes, respectively. Humidity was maintained at 70-80% with a 12-hour light and darkness photoperiod in all the rooms. Houseflies were collected from kitchens and living rooms of residential places within Nairobi, Kenya. The collected houseflies and fresh adult mosquitoes taken from rearing cages were anaesthetized using chloroform. This was done by enclosing the separate groups of the insects in enclosed chambers that contained open bottles of chloroform for six hours.

### Raman Spectroscopy Measurements

Raman spectra were acquired using a Technos® dispersive Raman microscopy system. The system had the following parameters: 532nm laser, 600 grooves per mm gratings, and ×10 infinity-corrected dry microscope objective with a Numerical Aperture of 0.25. Figure 1 shows a schematic of the instrument set-up with the sample (insect) placed on the X-Y translation stage. The insect sample was focused and viewed via the video monitor on the computer screen and simultaneously scanned by translating the X-Y stage to identify the region of interest. All the insects were scanned on the legs. Photons from the laser (indicated as green arrows pointing toward the sample) were delivered through the microscope objective to the sample, which was on a Raman-grade Calcium Fluoride microscope slide (Crystran Ltd, UK, Batch No. 60373). Raman and Rayleigh scattered photons (indicated as green and red arrows pointing away from the sample) were collected by the same microscope objective with an optical low pass filter blocking the latter. The Raman signal was collected via an optical fibre and spectrograph for digitization by a charge-coupled device (CCD) (cooled to -76°C) connected to a computer for display and storage. Wavenumber calibration was done by interpolating the laser line and the strong silicon Raman shift positioned at 520.5±4 cm^-1^.

**Figure 1.**
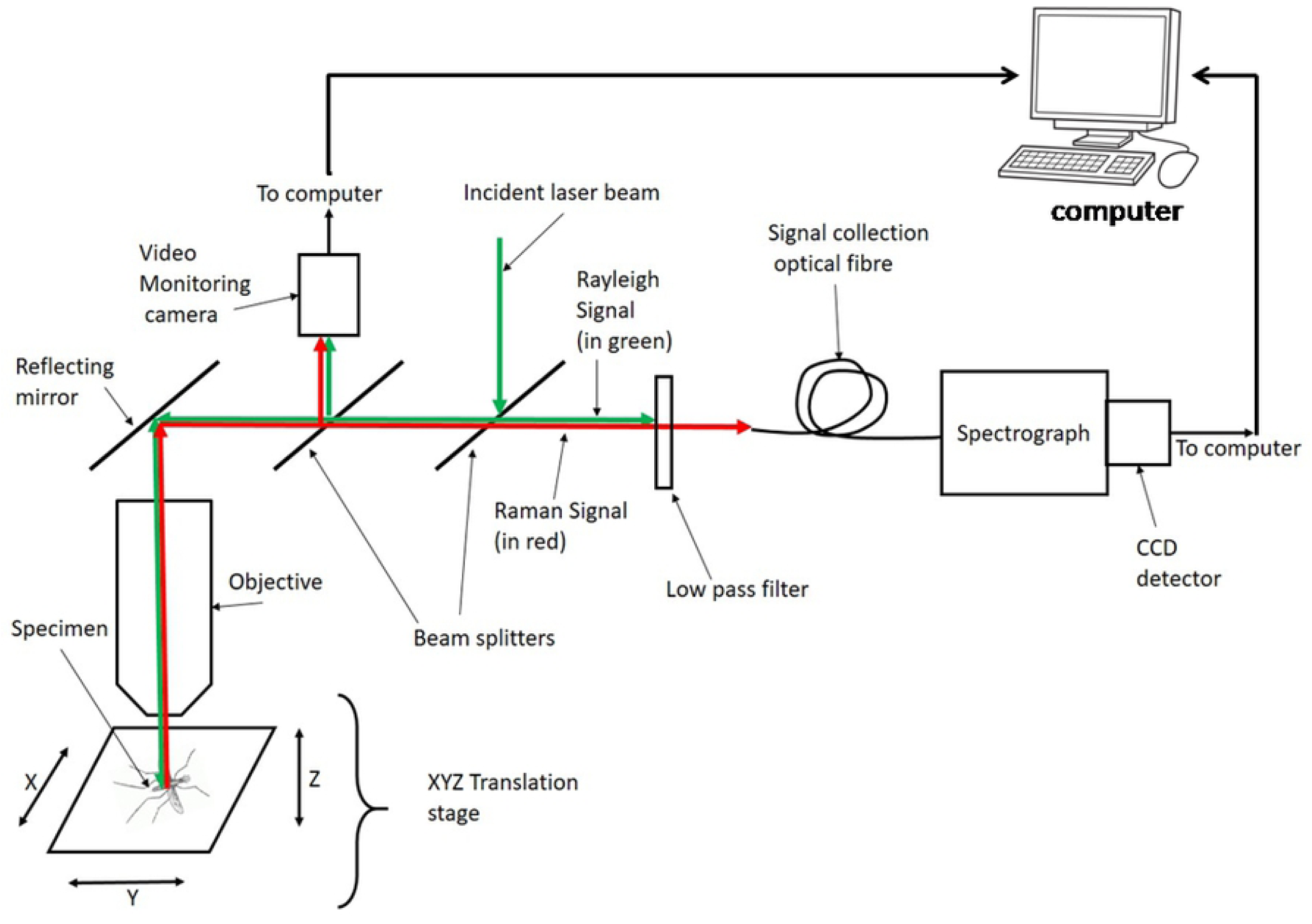
A schematic representation of a dispersive Raman microscope. The sample is placed on the X-Y stage. The red and green arrows indicate photon delivery routes.

### Data Pre-processing

Data processing was performed following previously published protocols by Ryabchykov (32) and Morais (33). Acquired data were smoothed by convolving each Raman spectrum with a Savitzky Golay digital filter of order 5 and frame length of 21 pixels. This was followed by a baseline correction procedure employing the Vancouver algorithm (34) with a 5^th^ order polynomial to subtract fluorescence. Vector normalization was applied to each Raman spectrum to account for intensity variation due to experimental factors such as changes in sample focus. All pre-processing procedures were done using scripts developed in Matlab^®^ 2018 software.

### Feature Selection and Feature Extraction

Data reduction consisted of two steps: feature selection followed by feature extraction. Feature selection involved the reduction of input variables by selecting a subset of Raman shifts that were considered most relevant for developing a classification model. Variance threshold method (VT) (35), a technique of feature selection drawn from a broad range of feature selection methods known as filter methods, was chosen due to its simplicity. The Raman data were stacked into a matrix *X* ∈R^m×n^ where m and n were the number of objects (the insects) and features (Raman shifts), respectively. The value of m was approximately 480 (consisting of approximately 120 insects of each of the categories: *Anopheles, Aedes, Culex* and houseflies). The Raman shifts (n=341) spanned wave number range 1000-2300 cm^-1^. Variance threshold (VT) was performed in Matlab^®^ 2018 software using

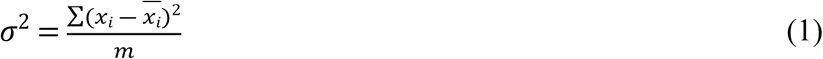

where σ^2^ is the variance of a feature and *x*_*i*_ is a vector containing *i*^th^ feature in the data matrix *X* ∈R^m×n^.

A total of 123 features with variance scores below a 0.0003 threshold value were excluded from the feature extraction step. The selected 218 features were subjected to feature extraction by performing Principal Component Analysis (PCA). PCA projected the selected feature data into a low dimensional subspace resulting in 24 orthogonal score variables that captured 94% of the information in selected features. All score features accounting for less than 0.5% variability were considered noise and excluded from the model. PCA numerical decomposition was performed in Matlab® 2018 software using

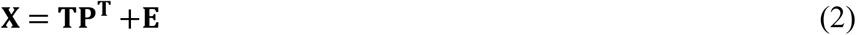

where **T** is a matrix containing scores, **P**^T^ is the transpose of the matrix containing loadings, and **E** is the residual.

### Classification Models

The extracted 24 principal components were subsequently subjected to Linear Discriminant Analysis (LDA) and Quadratic Discriminant Analysis (QDA) using OriginLab^®^ 2019 and Matlab^®^ 2018 software. Training and testing of both LDA and QDA were done by leave-one-out cross-validation. Training the models involved finding suitable decision boundaries between the classes. LDA and QDA, being generative models, relied on a full structured joint probability distribution over the training samples and labels. The basic assumption in the classification models was that the data followed a normal distribution. Therefore, each class label was fitted to a Gaussian distribution function using the calculated covariance matrices of the multivariate data during training and decision boundaries found based on the prior probability of each class. Prediction of test samples was achieved by evaluating each discriminant function, and the class label of the test sample was assigned to the highest-scoring function. The calculations were based on Tharwat’s guide (36), summarized as follows: The first 24 features of the score matrix, **T**, were used to find the decision boundaries. A decision boundary, S_12,_ between any two classes C_1_ and C_2_ with means μ_1_ and μ_2_ respectively, covariances Σ _1_ and Σ _2_ respectively, and probabilities P(C_1_) and P(C_2_) respectively, is defined, for QDA, as a quadratic function represented by

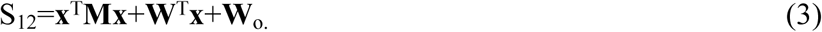

In equation 3, **x**^T^ is the transpose of **x**, the vector containing the classification features of each sample;

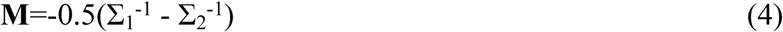

where Σ _1_ ^-1^ and Σ _2_ ^-1^ are inverses of the covariance matrices Σ _1_ and Σ _2,_ respectively;

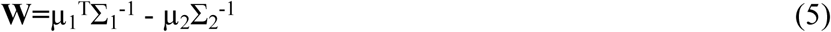

and

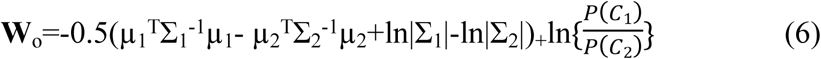

where |.| denotes the determinant of the enclosed matrix. For LDA, the decision boundary is evaluated by omitting the quadratic term, **x**^T^**Mx**, in equation 3.

### Performance Quality Metrics

Five quality metrics were calculated from the confusion matrices of the developed VT/PCA/LDA and VT/PCA/QDA models to evaluate their performance: Accuracy, Sensitivity, Specificity, F-score, and G-Score. Accuracy was defined as the percentage of correct classification; sensitivity the percentage of true positives that were classified correctly while specificity the percentage of true negatives that received the correct classification. F-Score accounted for the balance between Sensitivity and Specificity in the classes whereas G-Score accounted for the class sizes. The metrics were calculated using equations 7-11 (33)

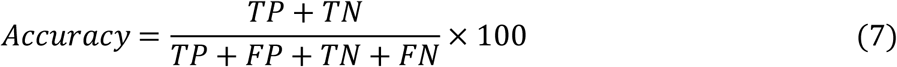

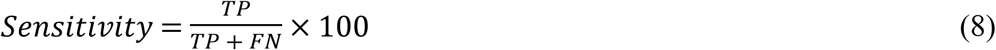

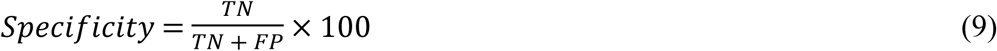

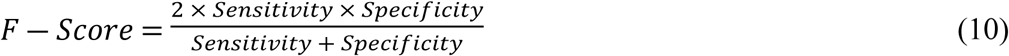

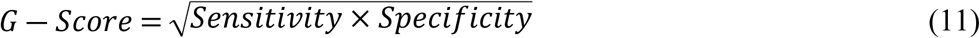

where TP, TN, FP, and FN represent True Positives, True Negatives, False Positives, and False Negatives respectively. Figure 2 summarizes the data analysis protocol followed in developing the models.

**Figure 2.**
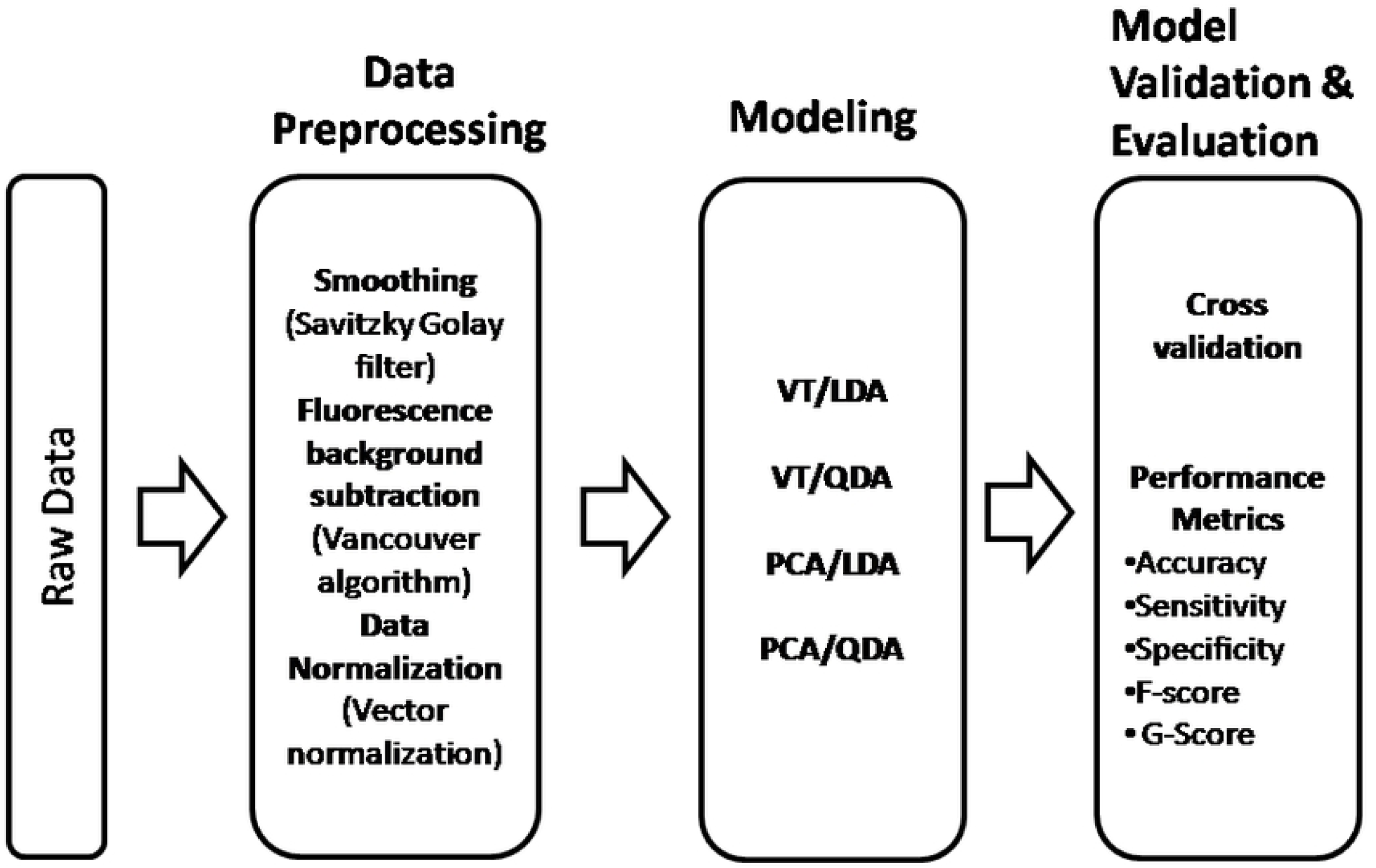
Data analysis pipeline: from the left, raw data is pre-processed, modelled, and finally, the models are validated and evaluated.

## RESULTS

The preprocessed Raman spectral data consisted of 341 features (wavenumber range 1000-2300 cm^-1^). The Raman shift range was chosen because it is known to correspond to the fingerprint region of many organic and biological molecules and hence was considered most suitable for classification. Figure 3 (a-d) shows Raman spectra of *Aedes aegypti, Anopheles gambiae, Culex quinquefasciatus* and *Musca domestica* (housefly). The spectra are dominated by broad peaks centred around 1400 cm^-1^, 1590 cm^-1^ and 2060 cm^-1^. These peaks are attributed to melanin (29,30,37–40).

**Figure 3.**
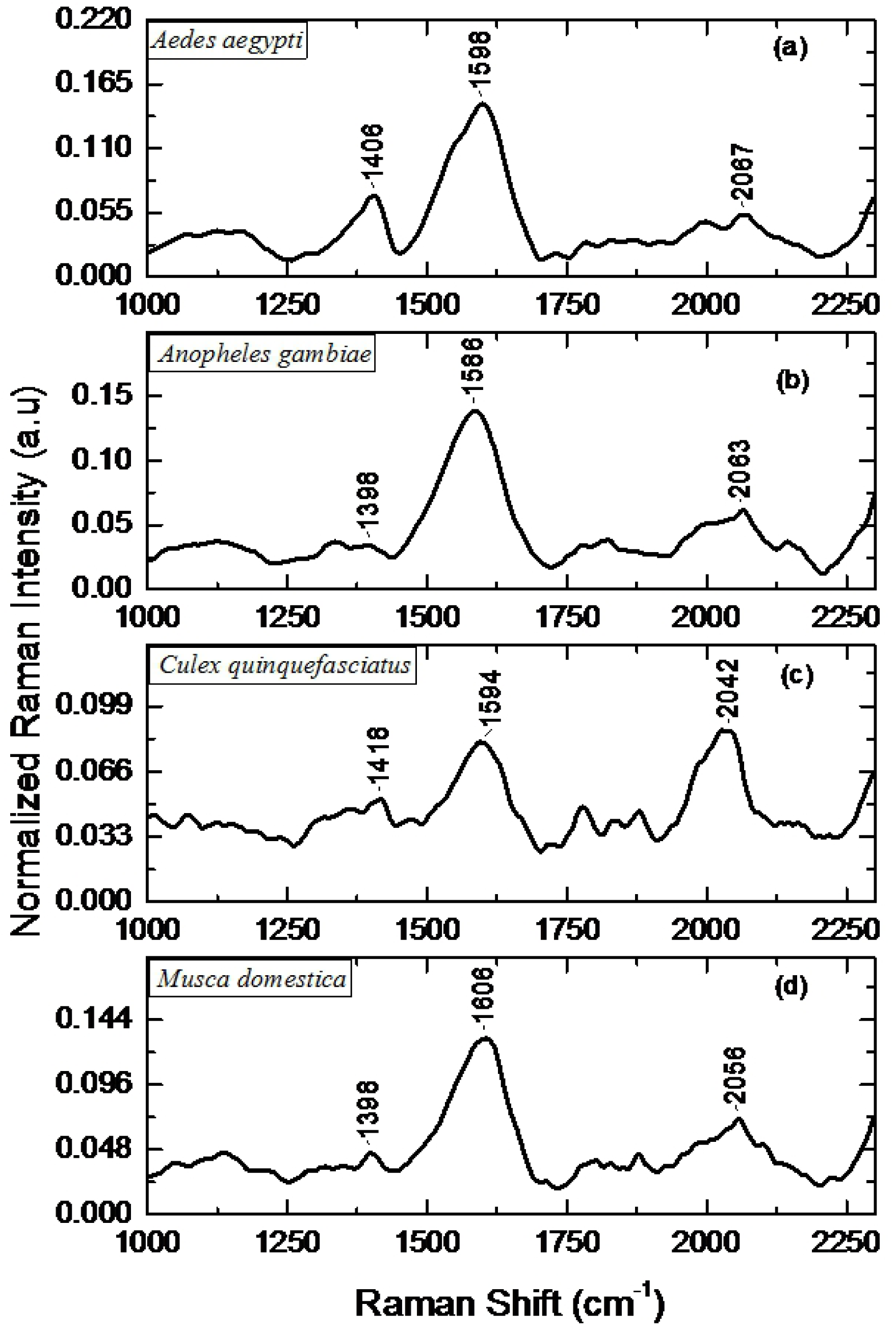
Preprocessed Raman spectra of (a) *Aedes aegypti*, (b) *Anopheles gambiae*, (c) *Culex quinquefasciatus* and (d) *Musca domestica*. The labelled peaks are those attributed to melanin pigment.

For discrimination purposes, features with large variance were needed since those with low variance have similar values across the four insect categories. Figure 4 shows the variance of each of the original features and the threshold value used to select the features. The greatest variance in the data set occurs around 1590 cm^-1^, followed by 2060 cm^-1^. Significant variances can also be seen on features around 1066 cm^-1^, 1315 cm^-1^, 1410 cm^-1^, 1462 cm^-1^, 1667 cm^-1^, 1766 cm^-1^, 2100 cm^-1^ and 2165 cm^-1^. A total of 123 features with variance scores below the 0.0003 threshold value were excluded from further processing. The value of 0.0003 was determined using an iterative process in which the threshold value that led to the best classification was identified. The large variance in the Raman bands centred around 1400 cm^-1^, 1590 cm^-1^, and 2060 cm^-1^ could, therefore, be ascribed to differences in quantities of the eumelanin and pheomelanin within the insect species.

**Figure 4.**
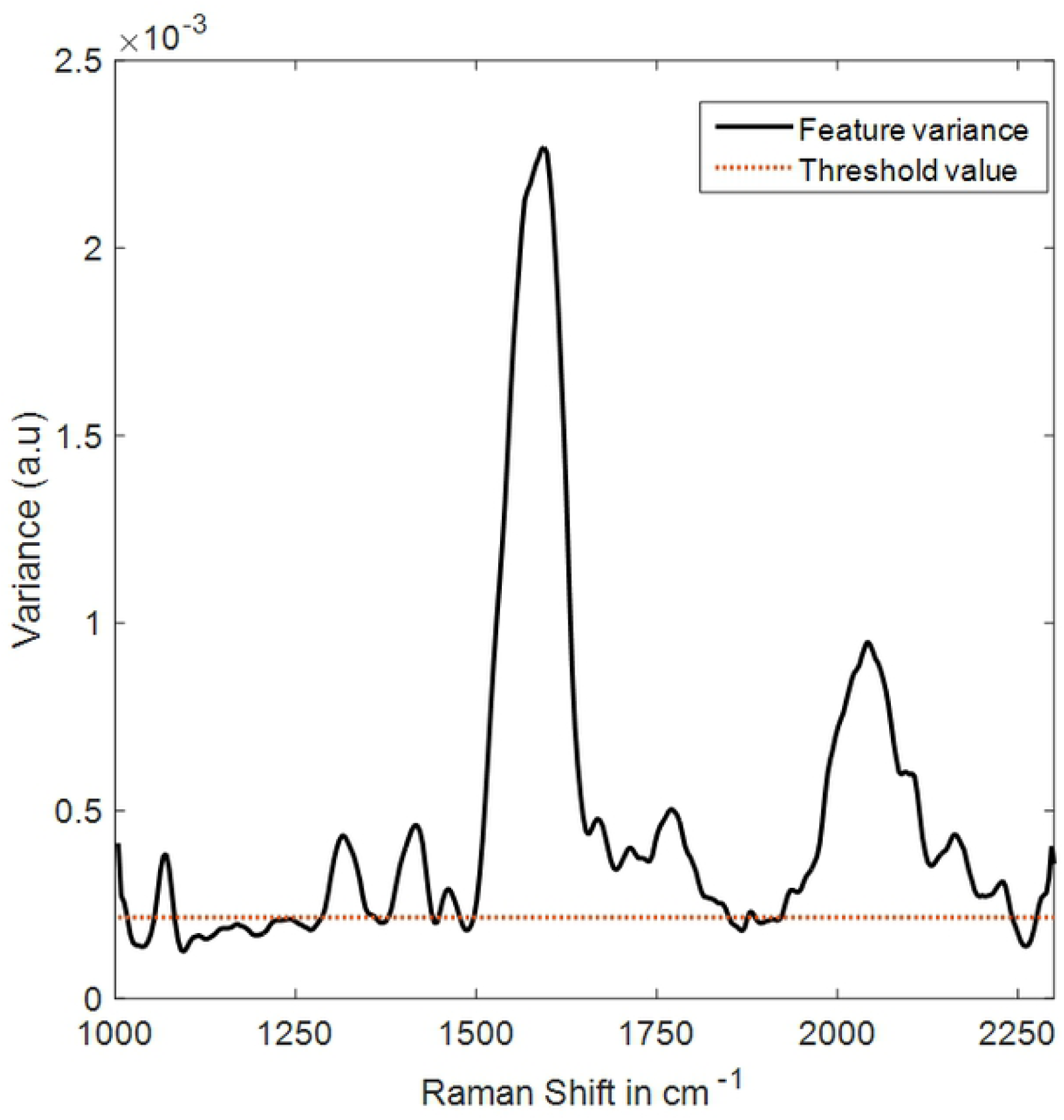
A plot of variance within each Raman band for all the four insect categories. The threshold value is indicated by the red dotted line.

In order to meet the dimensional requirements of Discriminant Analysis, the selected 218 features were compressed to 24 PC scores (represented by **T** in equation 2) which accounted for 94% variability. The 25th PC score and those beyond were each found to account for less than 0.5% variability and were considered to be noise.

The following two classification models were tested on the datasets: VT, then PCA, followed by LDA (VT/PCA/LDA) and VT, then PCA, followed by QDA (VT/PCA/QDA). VT/PCA/QDA performed better than VT/PCA/LDA achieving an overall accuracy of 94% against the 85% accuracy rate achieved by the latter. Table 1 summarizes the overall performance of the two models as assessed based on the five quality metrics: accuracy, sensitivity, specificity, F-Score and G-Score as defined by equations 7-11.

**Table 1.** Performance Quality Metrics for VT/PCA/LDA and VT/PCA/QDA Classification Models.

|  | Accuracy | Sensitivity | Specificity | F-Score | G-Score |
| --- | --- | --- | --- | --- | --- |
|  | (%) | (%) | (%) | (%) | (%) |
| VT/PCA/LDA | 85 | 69 | 90 | 77 | 78 |
| VT/PCA/QDA | 94 | 87 | 96 | 91 | 91 |

The numerical figures presented in Table 1 are averaged values of the figures of merit as calculated from the confusion matrices of the VT/PCA/LDA and VT/PCA/QDA classifiers. Tables 2 and 3 present the confusion matrices that resulted from the VT/PCA/LDA and VT/PCA/QDA classifiers. The numerical figures in these matrices are the actual numbers of insects used in the cross-validation of the models. In examining each confusion matrix, it became clear that the two classifier models performed much better in distinguishing Anopheline mosquitoes (represented by *Anopheles gambiae)* from Culicines (represented by *Aedes aegypti, Culex quinquefasciatus)*. For instance, the models were more ‘confused’ in distinguishing between *Aedes* versus *Culex* than *Aedes* versus *Anopheles* for both models. In Table 3, 30 *Culex* mosquitoes were identified as *Aedes* as opposed to zero *Anopheles* confused for *Aedes*. A similar trend is observed in Table 2, in which 34 *Culex* mosquitoes were classified as *Aedes* as opposed to 4 *Anopheles* identified as *Aedes*. Thus, VT/PCA/QDA performed better than VT/PCA/LDA in classifying Culicines from Anophelines. However, both models were more likely to classify houseflies as Culicine mosquitoes.

**Table 2.** Confusion Matrix of VT/PCA/LDA Classifier.

|  |  | Predicted Class |  |  |  |
| --- | --- | --- | --- | --- | --- |
|  |  | *Aedes* | *Anopheles* | *Culex* | Housefly |
| Actual Class | *Aedes* | 132 | 8 | 22 | 6 |
|  | *Anopheles* | 4 | 92 | 12 | 12 |
|  | *Culex* | 34 | 6 | 70 | 18 |
|  | Housefly | 27 | 8 | 7 | 79 |

**Table 3.** Confusion Matrix of VT/PCA/QDA Classifier.

|  |  | Predicted Class |  |  |  |
| --- | --- | --- | --- | --- | --- |
|  |  | *Aedes* | *Anopheles* | *Culex* | Housefly |
| Actual Class | *Aedes* | 152 | 0 | 14 | 2 |
|  | *Anopheles* | 0 | 120 | 0 | 0 |
|  | *Culex* | 30 | 0 | 92 | 6 |
|  | Housefly | 7 | 0 | 8 | 106 |

To understand the differences between the decision boundaries established by the two classifier models during training, the discriminant scores, calculated using equation 3, were plotted to show the decision borders. Figure 5 shows the evaluation of the linear term of equation 3, which represents hyperplane decision boundaries between the four categories of insects. Each of the boundaries is represented by points where the equation evaluates to zero between any two categories of insects, with the separated categories assuming either positive or negative values. It can be observed that the decision boundaries created by VT/PCA/LDA are plagued with lots of overlap, especially for categories that exhibit large variances. The variances of the various classes can be visualized by how close the members of each class are to each other in the discriminant score plots. For instance, in Figure 5 (a), the *Anopheles* mosquitoes are well identified by negative discriminant scores, while the *Aedes* assume positive scores. However, due to large variation within the *Aedes* group, a portion of *Aedes* were discriminated as *Anopheles* by assuming negative discriminant score values. This trend is repeated in Figures 5 (b), (c) and (f). In Figures 5 (d) and (e), it is equally noted that the *Anopheles* are well discriminated with positive values of the discriminant scores, but the large variation within *Culex* and houseflies results in poor decision boundaries.

**Figure 5.**
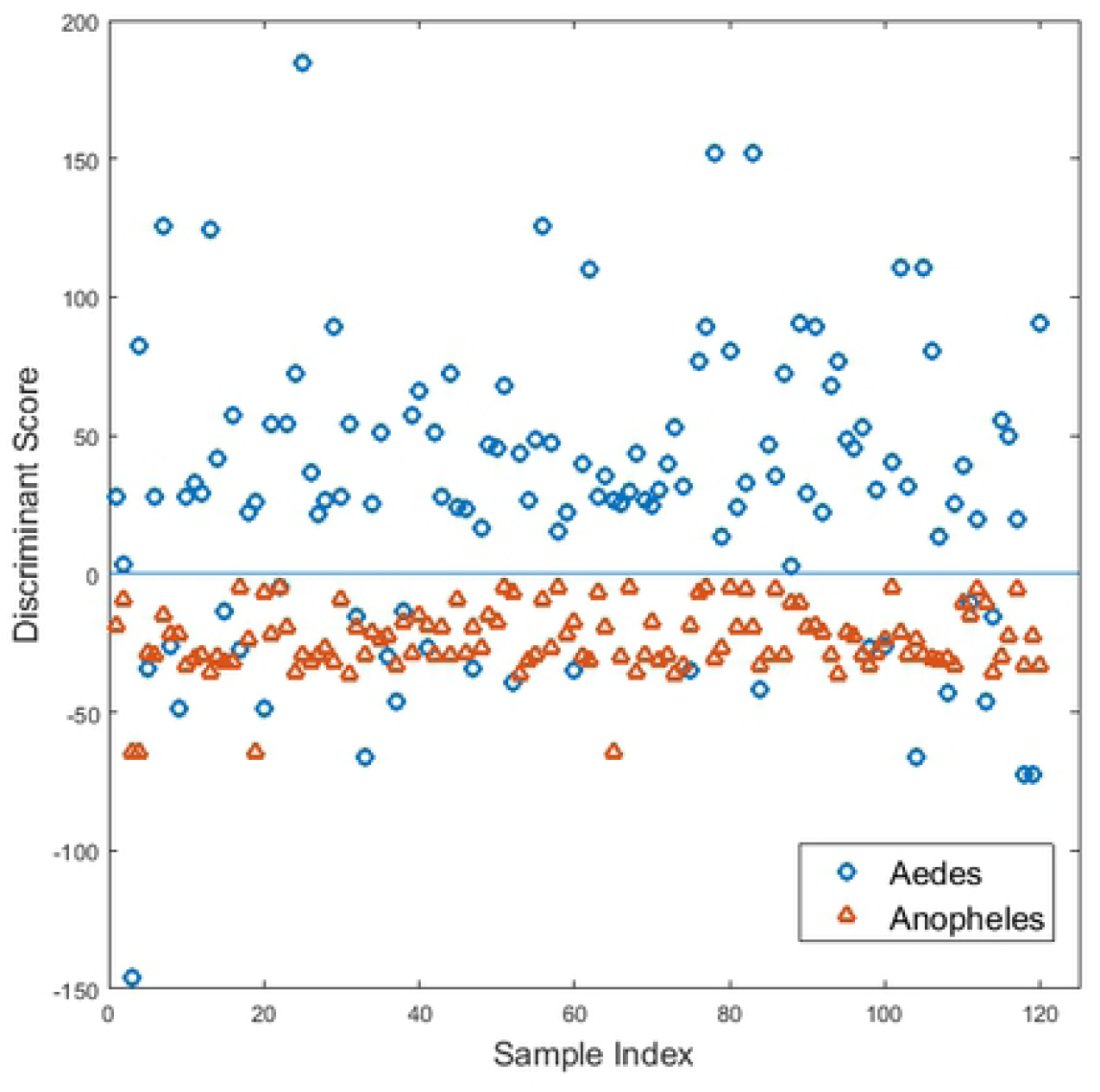
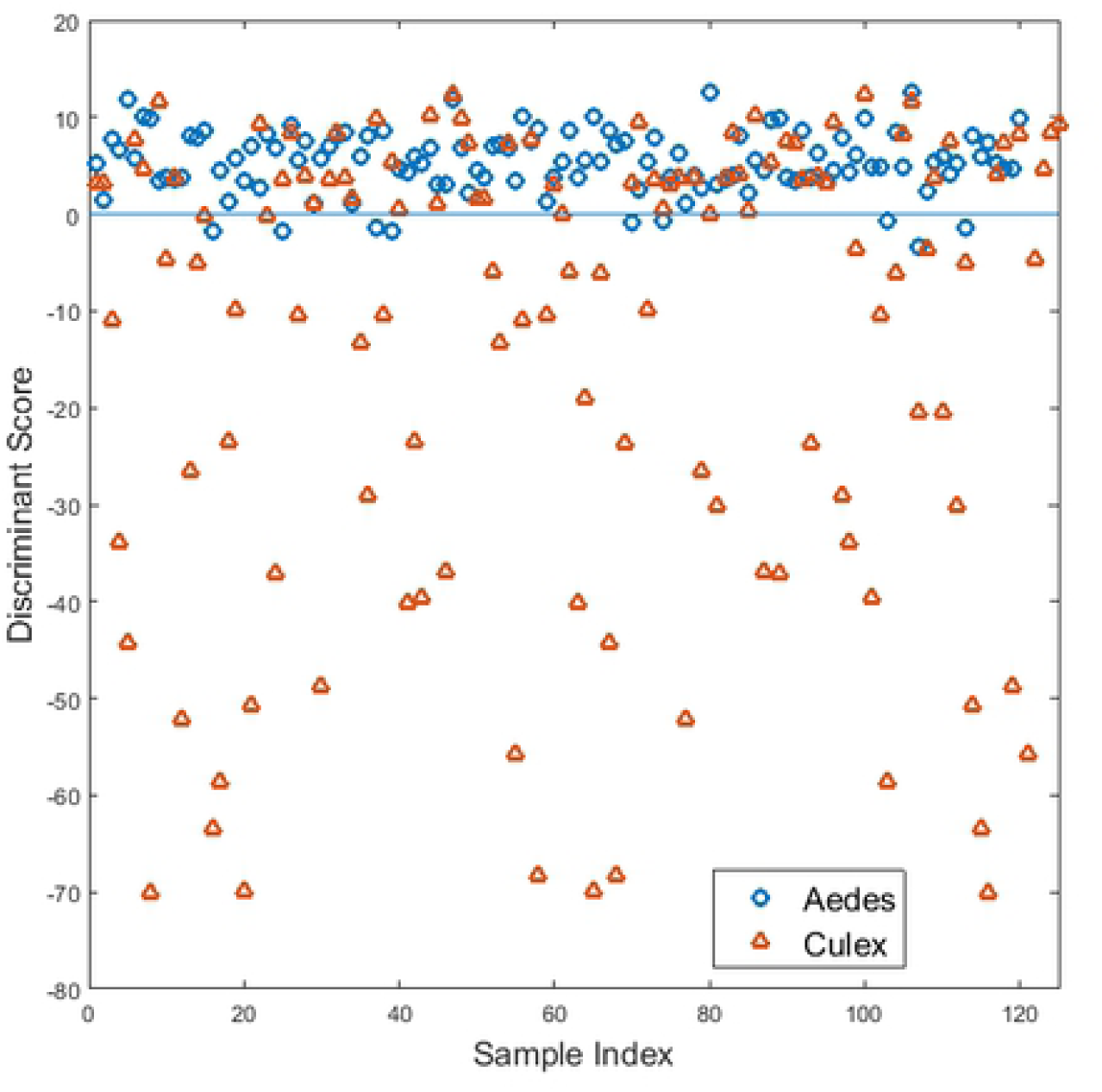
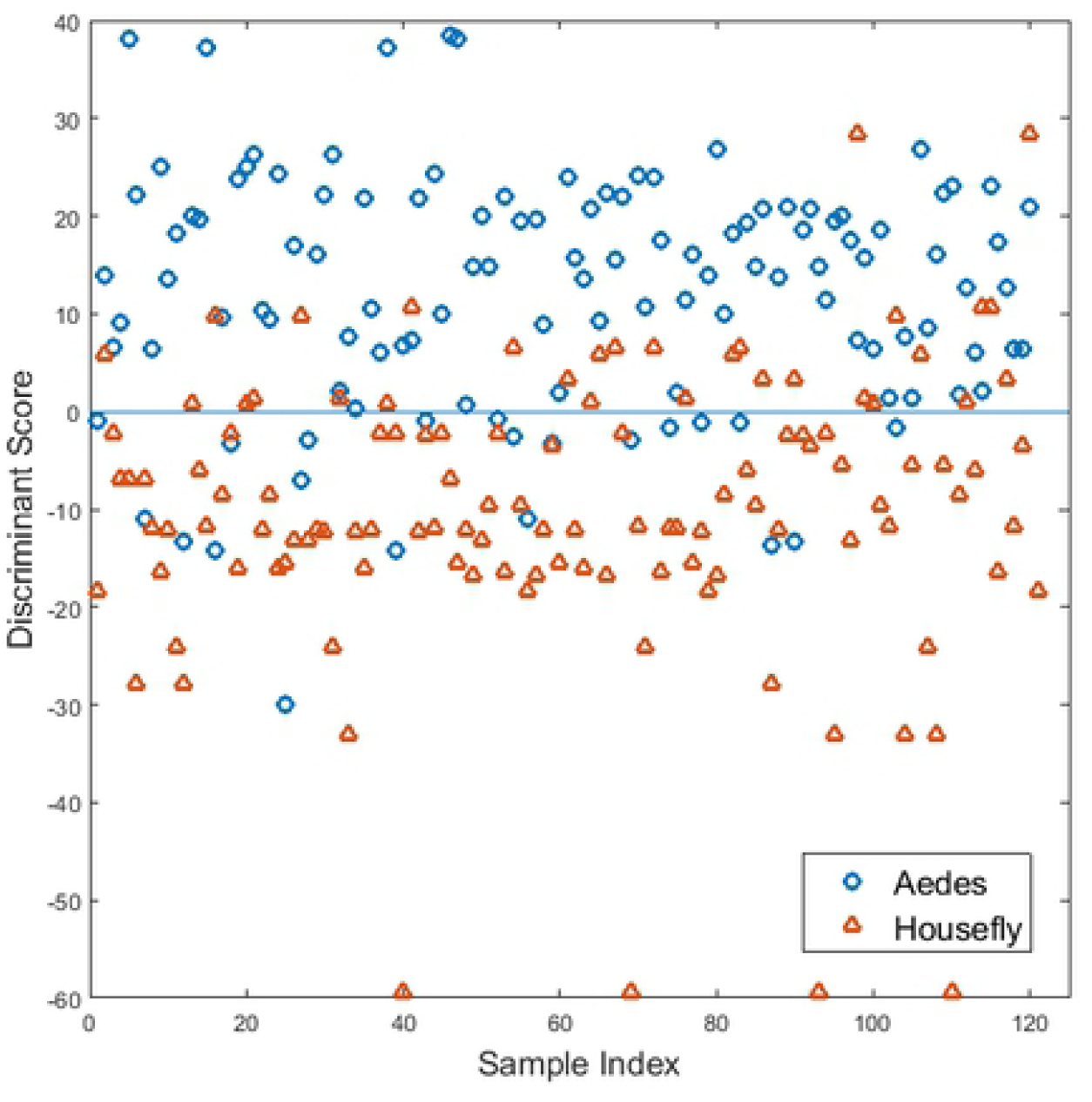
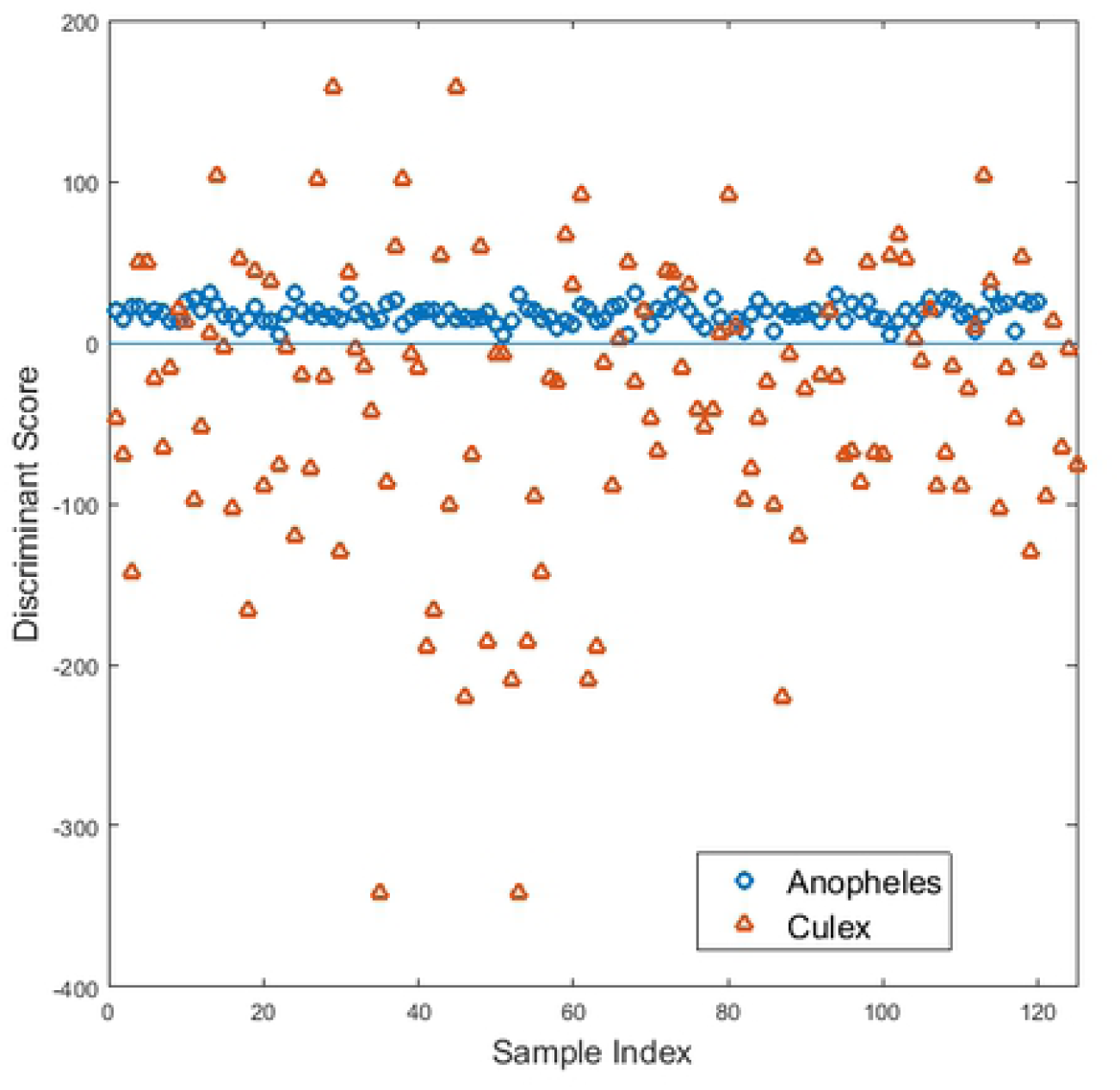
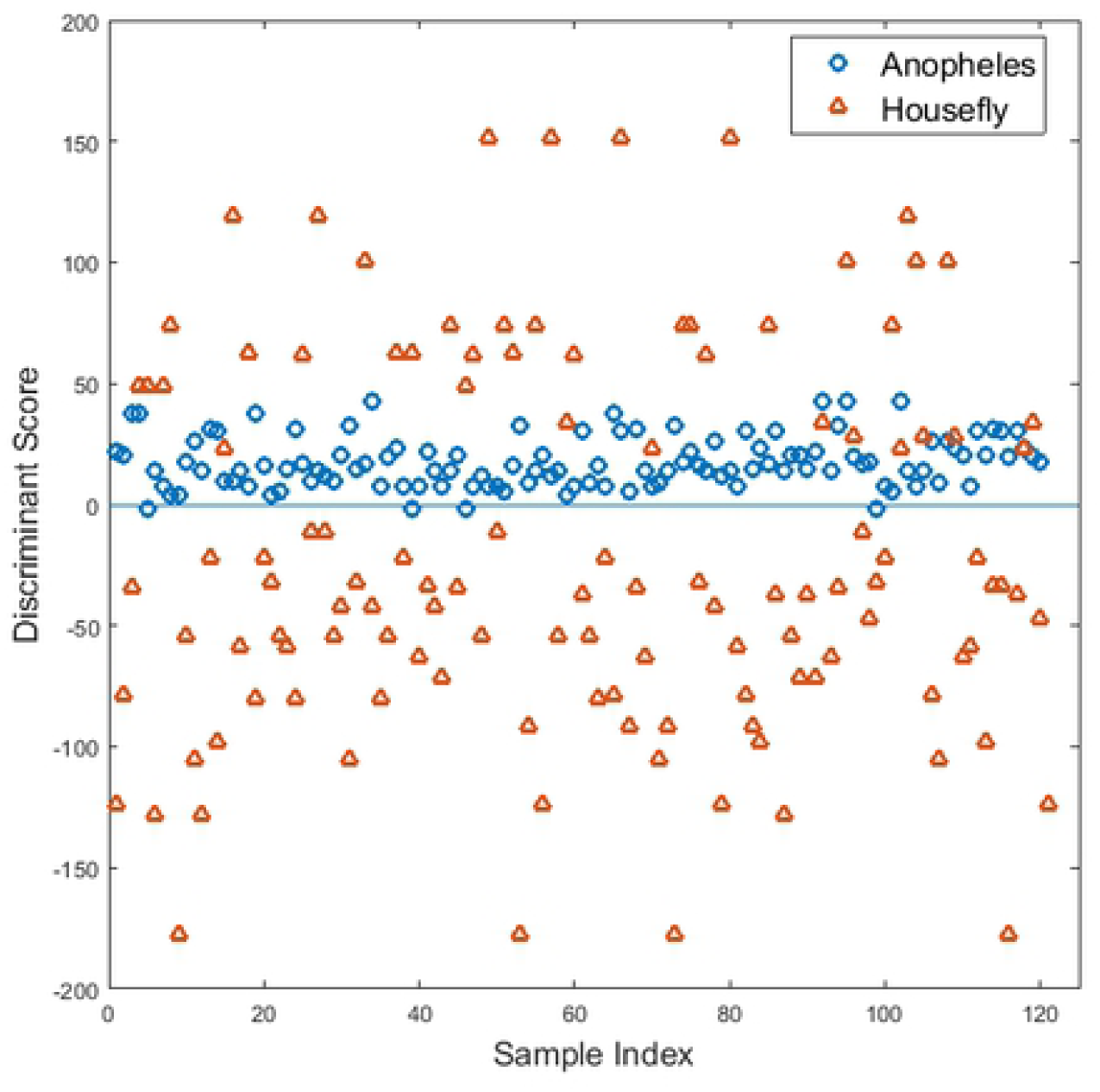
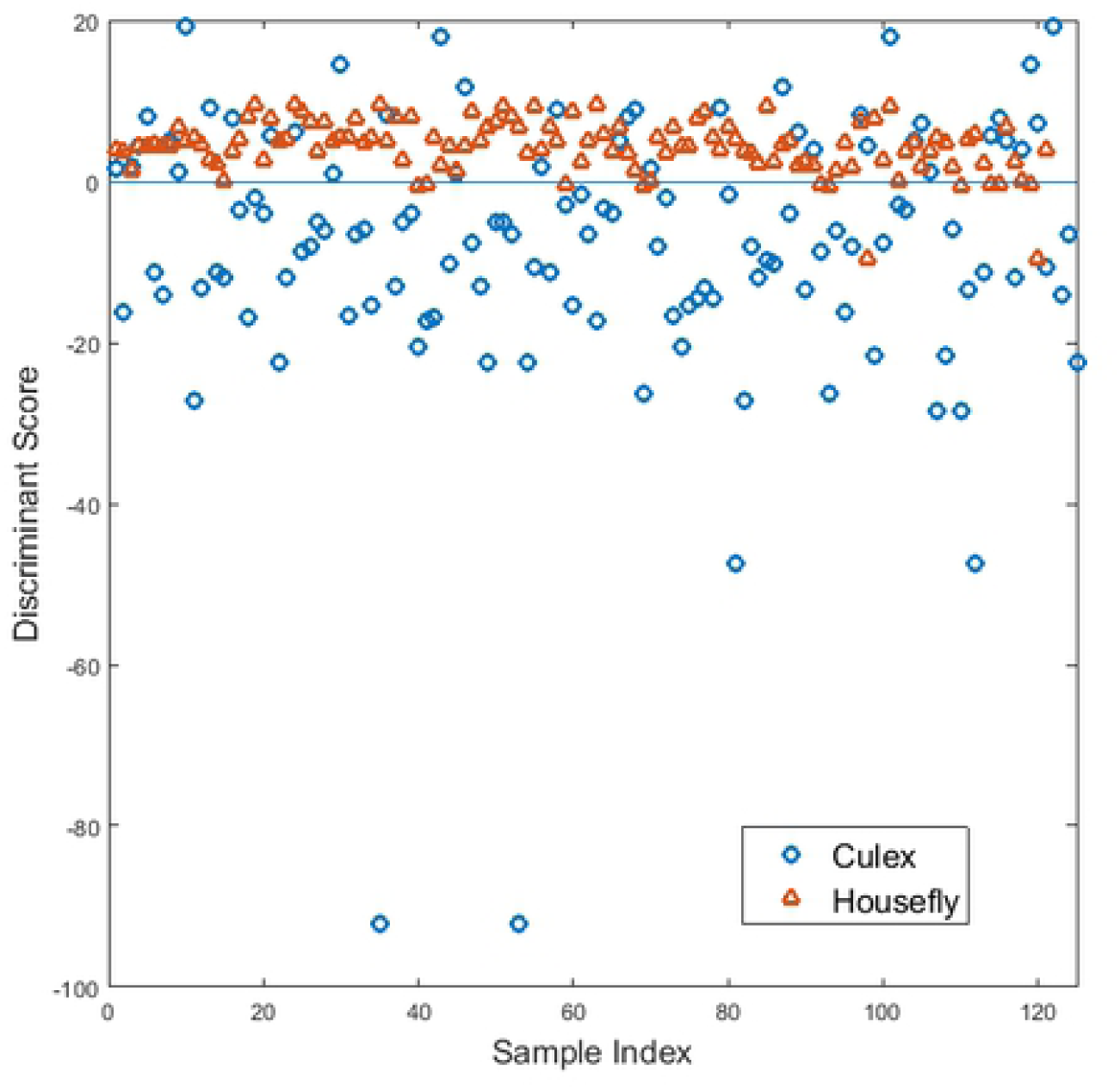
Decision boundaries as evaluated by VT/PCA/LDA. Each class takes either a positive or negative value Discriminant Score. (a) *Aedes* versus *Anopheles*, (b) *Aedes* versus *Culex*, (c) *Aedes* vs Housefly, (d) *Anopheles* versus *Culex*, (e) *Anopheles* versus Housefly, and (f) *Culex* versus Housefly.

VT/PCA/QDA model decision boundaries were evaluated using the quadratic and linear terms of equation 3. Figure 6 (a-f) shows the discriminant scores for the various group combinations. As explained earlier, the decision boundaries are represented by points where the equation evaluates to zero. The problem of different variances within the insect categories observed in Figure 5 (a-f) is also in Figure 6 (a-f). However, VT/PCA/QDA was able to discriminate well the various categories of insects during training by evaluating the non-linear portion of equation 3. The shortcoming of LDA is that it assumes that all the classes have a pooled covariance matrix which results in a linear decision boundary. QDA, on the other hand, considers the different covariance matrices for each class, which results in a quadratic decision boundary.

**Figure 6.**
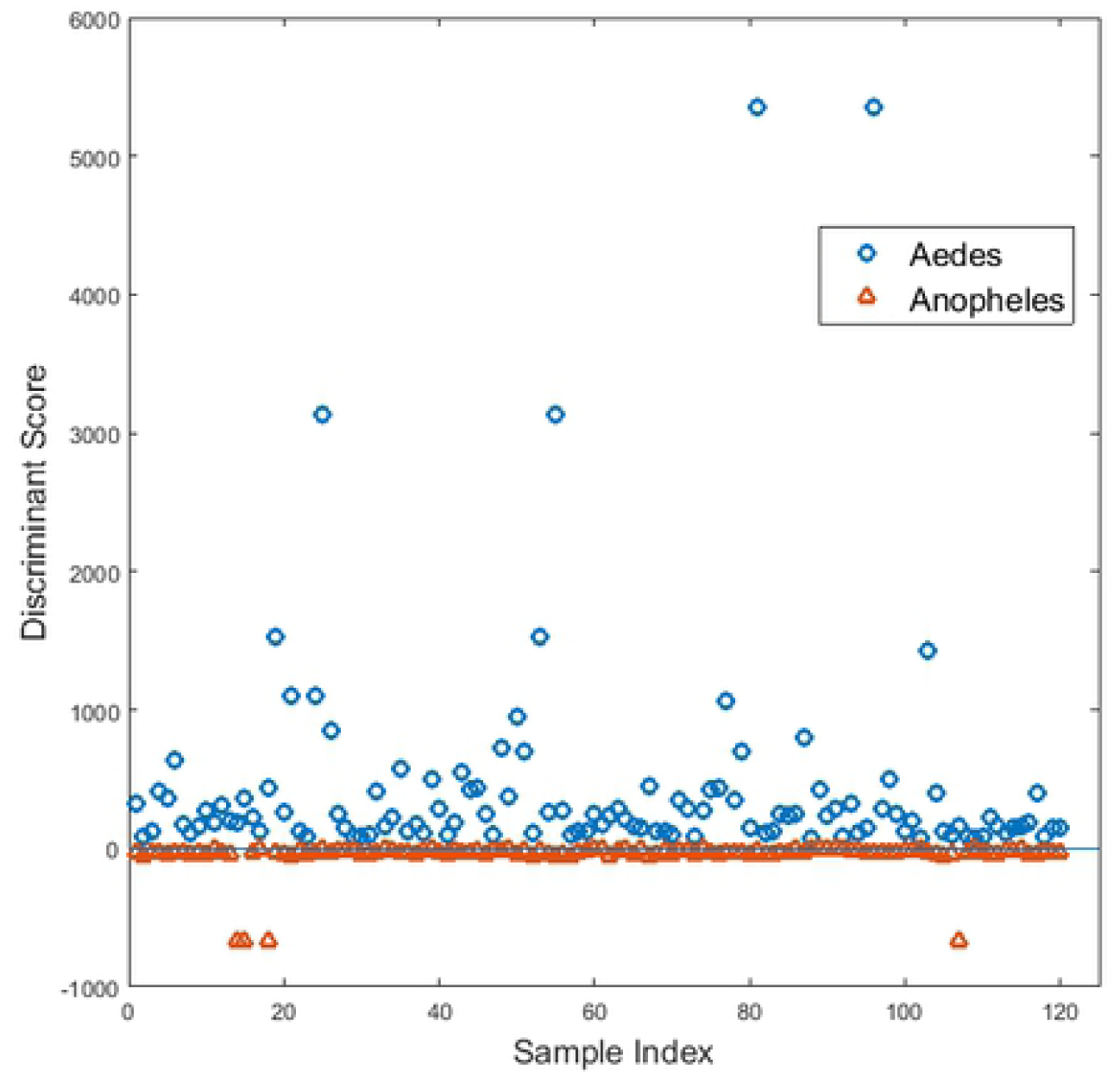
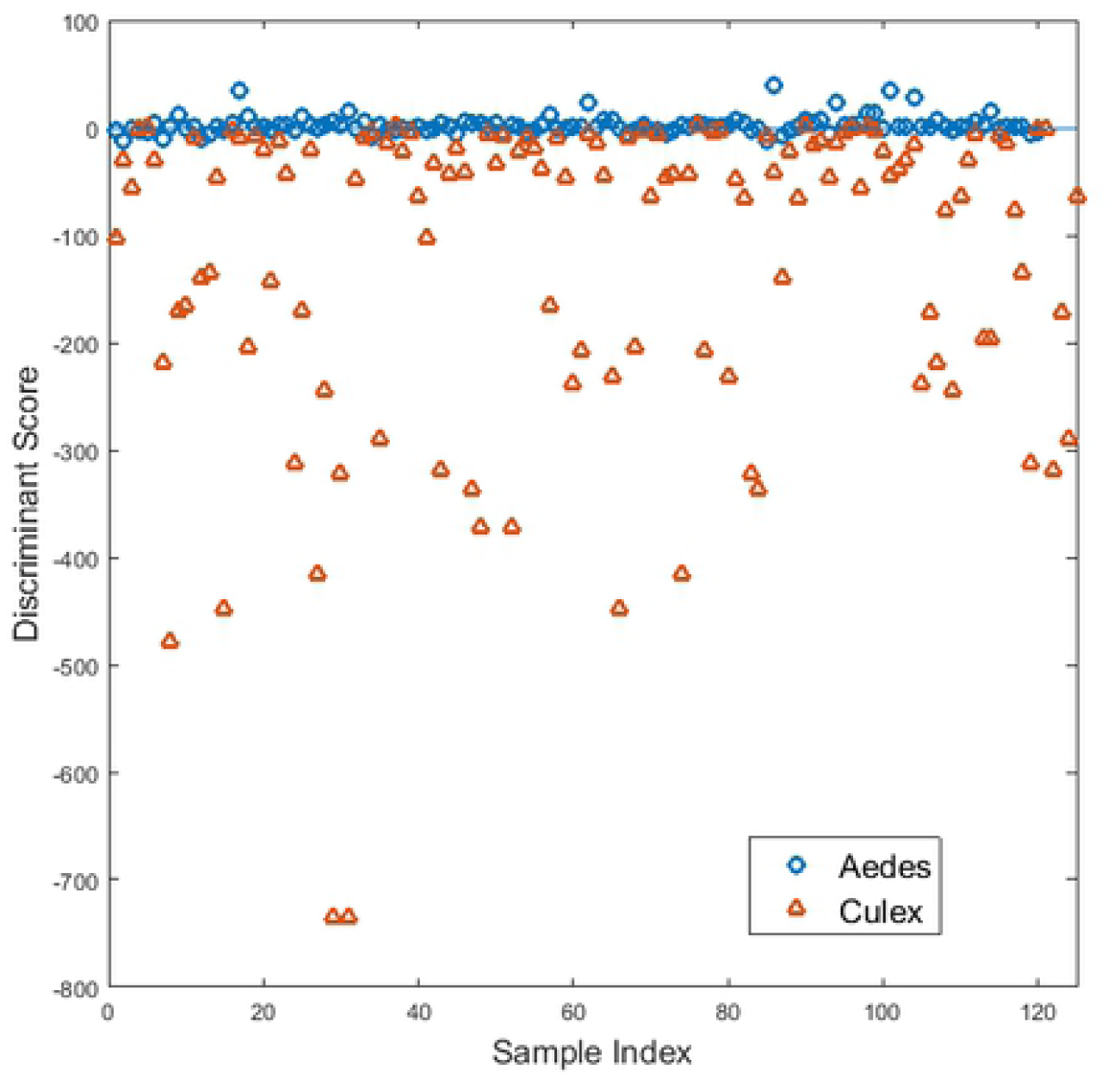
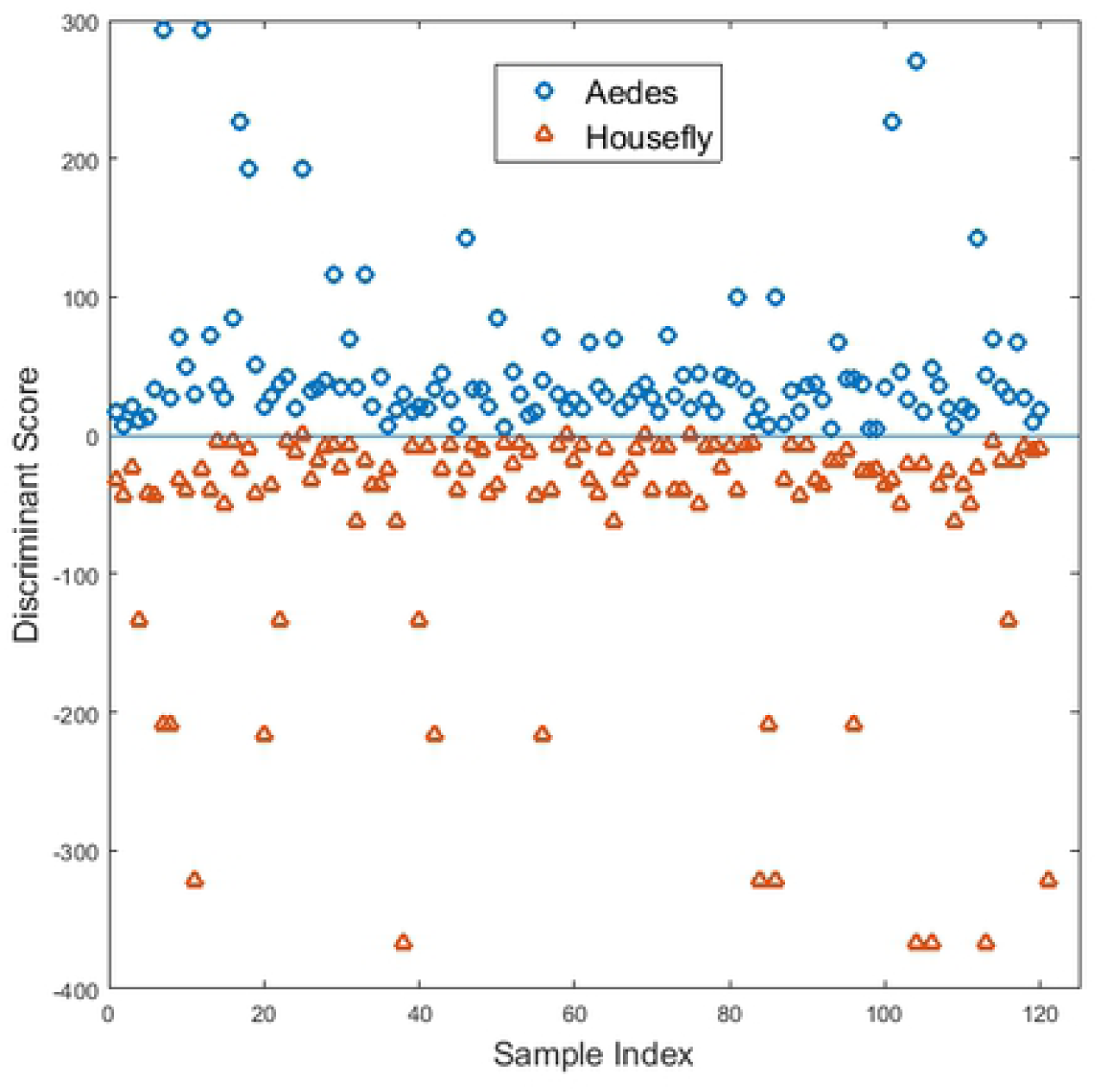
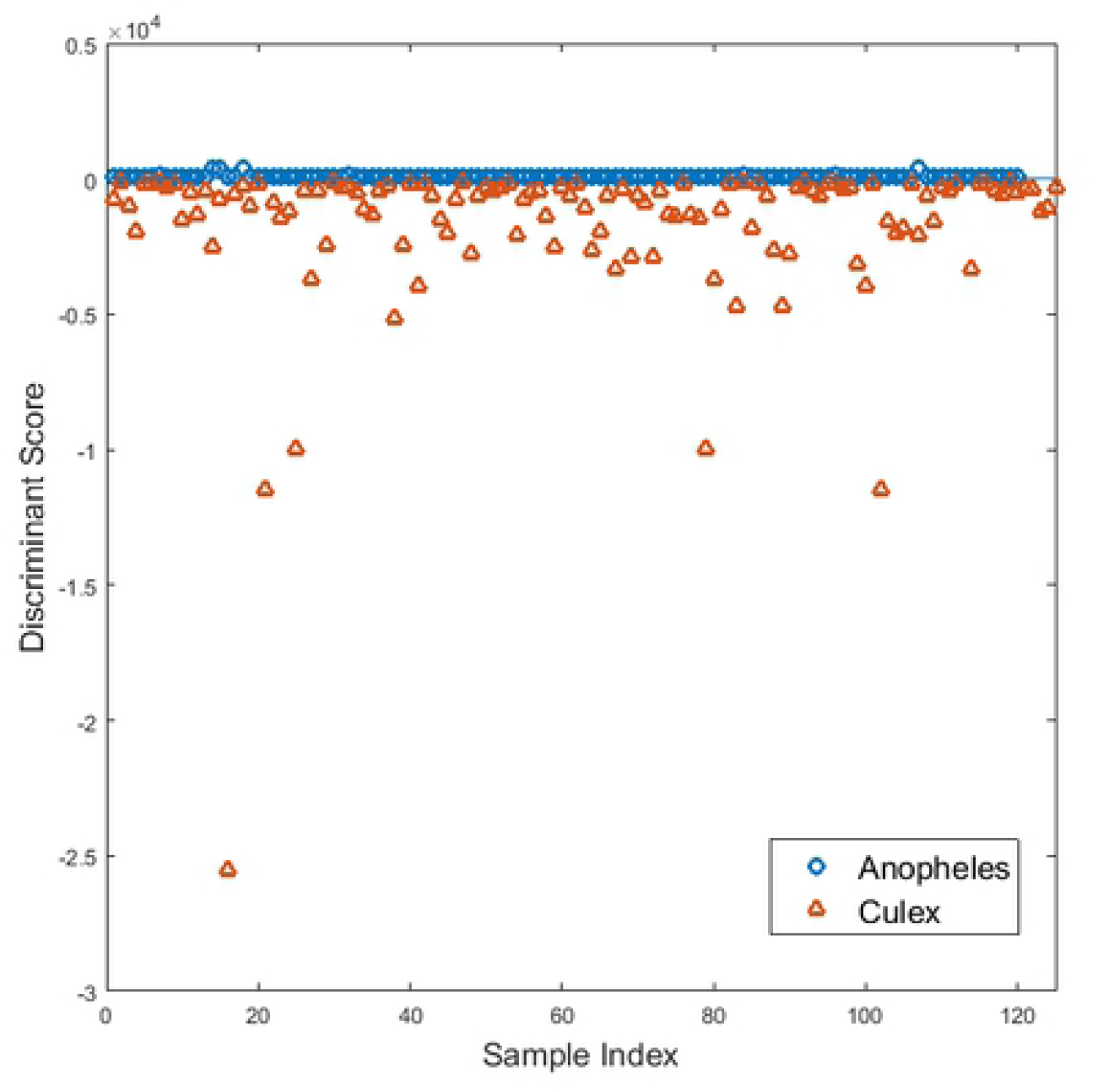
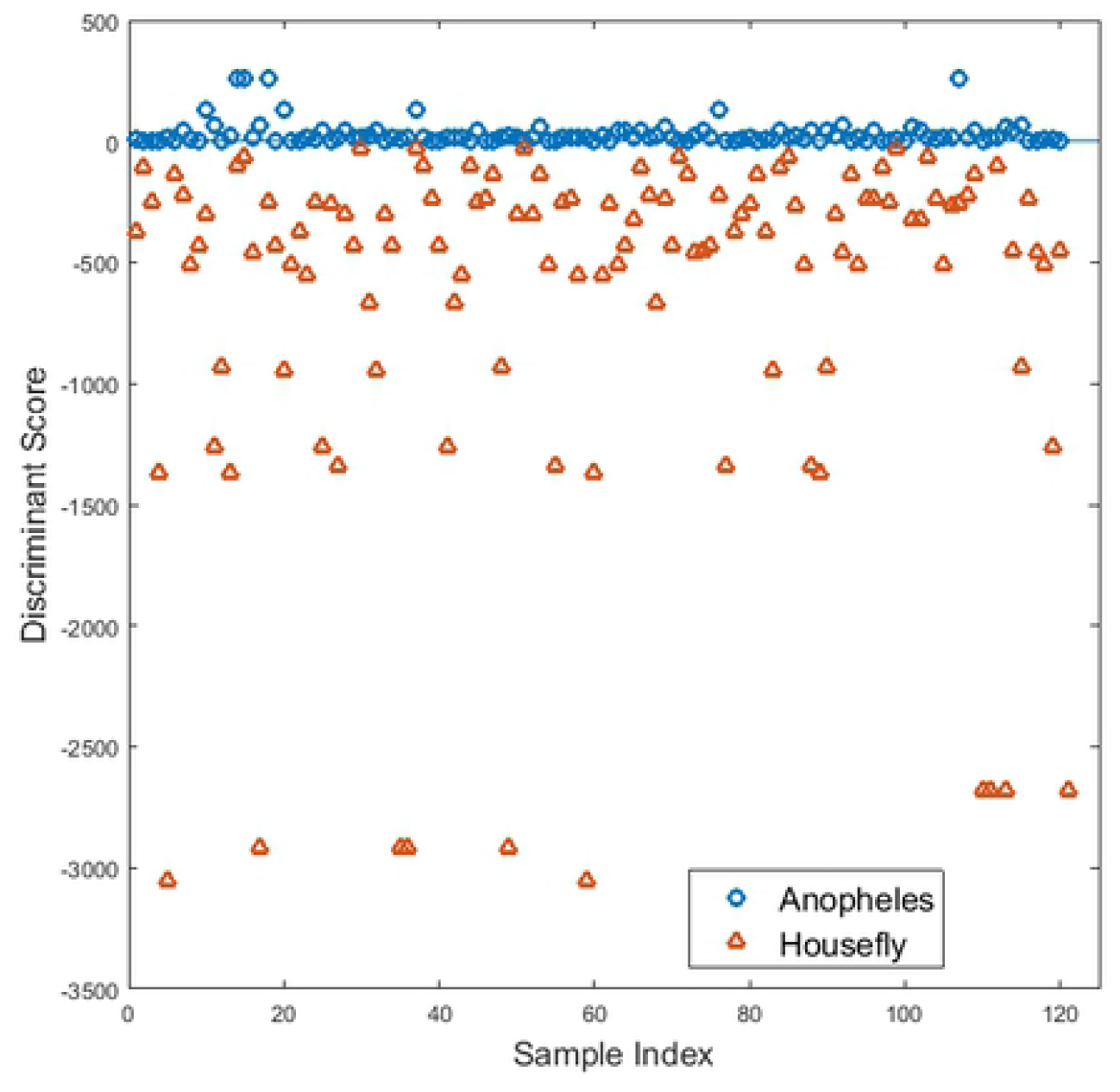
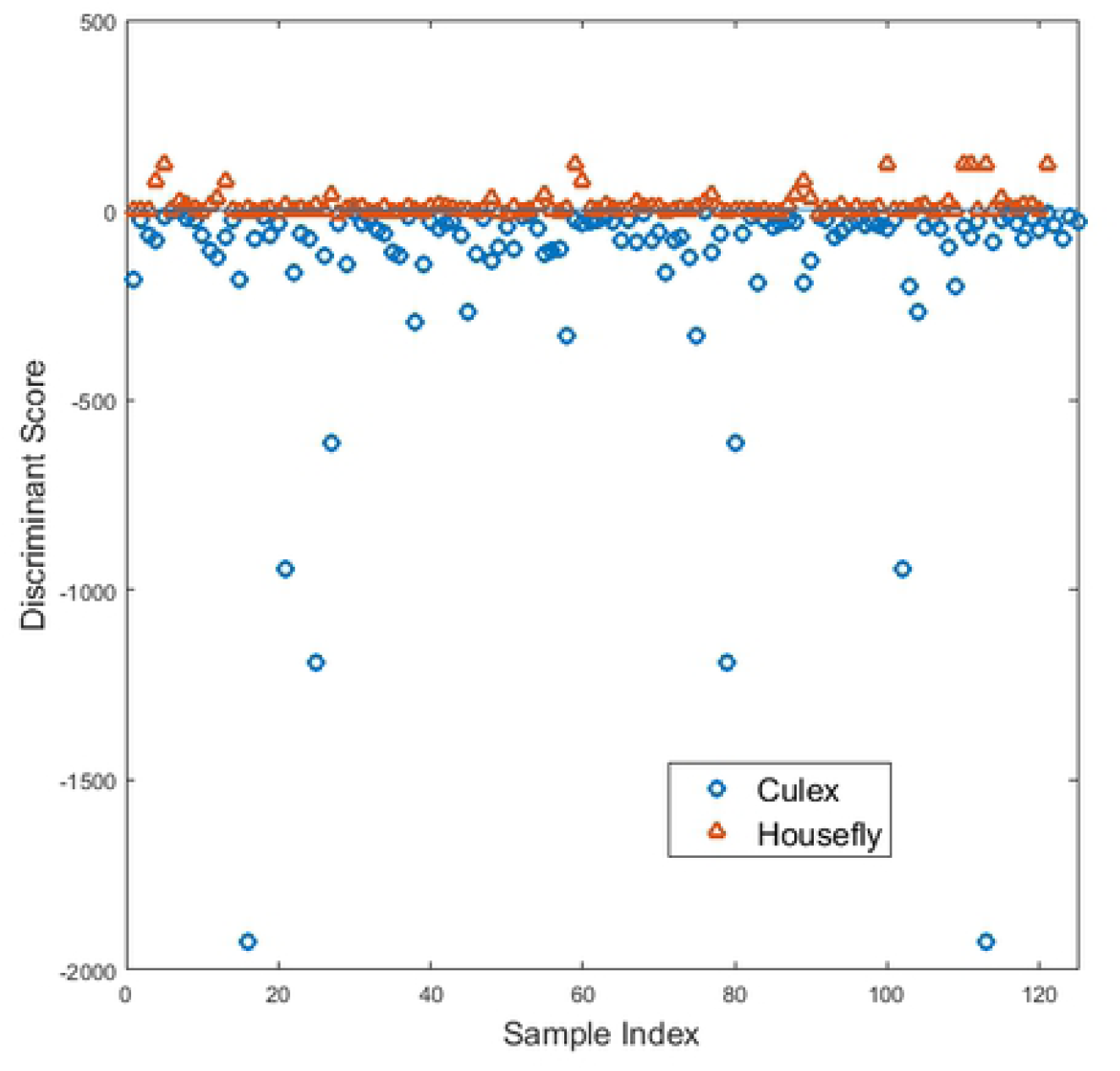
Decision boundaries as evaluated by VT/PCA/QDA. Each class takes either a positive or negative value Discriminant Score. (a) *Aedes* versus *Anopheles*, (b) *Aedes* versus *Culex*, (c) *Aedes* vs Housefly, (d) *Anopheles* versus *Culex*, (e) *Anopheles* versus Housefly, and (f) *Culex* versus Housefly.

## DISCUSSION

We have presented a proof of concept that three medically important mosquito species –*Aedes aegypti, Anopheles gambiae*, and *Culex quinquefasciatus* –can be classified based on Raman signals obtained from the surface of their leg cuticles. Raman signals were carefully extracted from the measured raw spectra using computer algorithms. The algorithms suppressed the auto-fluorescence background inherent in biological specimens, filtered out the noise and normalised the resulting Raman signals to provide a common basis for comparing samples. The Raman signals acquired from the three mosquito species in this study were dominated by broad peaks centred around 1400 cm^-1^, 1590 cm^-1^and 2060 cm^-1^. These peaks are due to melanin, a pigment found within insect cuticles (29,30,37–40). The significant spectral variance observed across the samples in these three spectral ranges provided important classification features for model development and highlights melanin’s potential as a biomarker for mosquito taxonomy. Two models, VT/PCA/LDA and VT/PCA/QDA, achieved 85% and 94% accuracy, respectively. This performance can be considered sufficient for the cost-effective screening of large numbers of mosquito samples usually collected in mosquito surveillance programs, field studies, and cases where samples have lost morphological features during storage.

Our demonstration that melanin can be used for taxonomy is a significant shift from previous spectroscopic classification work on mosquitoes which relied on signatures of cuticular hydrocarbons (24–26) and proteins (17–19). The prevalence of Raman peaks associated with melanin was a surprise since Raman spectroscopy had been expected to reveal signatures associated with proteins and lipids. Naturally, the variance at 1667 cm^-1^ was attributed to the Amide I band due to proteins (41) while those at 1066 cm^-1^, 1315 cm^-1^ and 1462 cm^-1^ were attributed to hydrocarbon chains of lipids (42). However, the most significant variance was attributed to eumelanin at 1590 cm^-1^ and pheomelanin at 2060 cm^-1^. Melanins have traditionally been considered difficult to extract for chemical analysis due to their low solubility (43); hence they have not been explored for insect classification. The ability of Raman spectroscopy to detect melanin signatures in mosquito cuticles makes melanin a potential biomarker for taxonomy. Melanin is the primary pigment responsible for colouration in animals and insects. In the latter, it is employed ingeniously for exoskeletal pigmentation, cuticular hardening, wound healing (44), and protection from solar radiation (45), among other innate immune responses. There are two main categories of melanin pigment - eumelanin and pheomelanin (46). Eumelanin is primarily responsible for dark colours, from brown to black, whereas pheomelanin produces yellowish or reddish colours.

In Figure 3, the peaks occurring around 1400 cm^-1^ and 1590 cm^-1^ were attributed to eumelanin, while the broad peak around 2060 cm^-1^ was attributed to pheomelanin (30,39,40). A closer look at Figure 3 (a) reveals that the eumelanin peak at 1598 cm^-1^ is much stronger when compared to the pheomelanin peak at 2067 cm^-1^. The 1406 cm^-1^peak is also well defined. This spectrum was taken from the dark portion of *Aedes aegypti* (the white part of the leg did not yield any significant peaks) and confirmed the black colouration of this insect species, an indication of the prevalence of eumelanin. In mosquito identification, colour generally plays a minor role, with descriptions of colour features limited to terms such as ‘ornamentation’ or ‘dark spots’, as is the case for *Aedes aegypti*. It is known that colour in insects emanates from pigments, mainly melanin, and structures that enhance visual appearance (47,48). Colour perception is very subjective in humans, but when measurements are taken using a spectral device like a Raman microscope, details that are not discernible by the naked eye are usually revealed. Figures 3 (b) and 3 (c) show spectra obtained from *Anopheles gambiae* and *Culex quinquefasciatus*, respectively. Although the two mosquitoes are generally described as ‘brown’ with minor variations described as ‘pale spots of yellow, white or cream scales’ in *Anopheles gambiae*, the spectra reveal that their eumelanin-pheomelanin combination is completely different and therefore useful for their discrimination. In Figure 3 (b), the peak at 1586 cm^-1^ (eumelanin) is much stronger than that at 2063 cm^-1^ (pheomelanin), whereas, in Figure 3 (c), the two peaks at 1594 cm^-1^ (eumelanin) and 2042 cm^-1^ (pheomelanin) are almost equal in strength. It should also be noted that there are variations in the positions and widths of the eumelanin and pheomelanin peaks across all the four groups (Figure 3 (a-d)) which could be attributed to the chemical environment (49) or the presence of other chemical compounds within the insect cuticle.

The confusion matrices (Tables 2 and 3) reveal how the VT/PCA/LDA and VT/PCA/QDA models responded when presented with each insect to classify. Overall, the models performed well in distinguishing between Anophelines (*Anopheles gambiae*) from Culicines (*Aedes aegypti* and *Culex quinquefasciatus*). However, the tendency of both models to misclassify houseflies for Culicine mosquitoes (*Aedes aegypti* and *Culex quinquefasciatus*) was puzzling. Probably this could be an indicator of yet to be known biochemical similarities between Culicine mosquitoes and houseflies, but which are not in Anopheline mosquitoes. We speculate that since flies have a common phylogeny (50), their underlying genomes may explain this unexpected similarity. Different species have conserved or modified their gene sequences as they evolved. The genome sizes of *Aedes aegypti, Anopheles gambiae, Culex quinquefasciatus* and *Musca domestica* are known to be 1.38 Giga bases (Gb), 278 Mega bases (Mb), 579 Mb, and 691 Mb, respectively (51–54). Therefore, from the genome sizes, *Aedes aegypti, Culex quinquefasciatus* and *Musca domestica* may have conserved some orthologs that were deleted in *Anopheles gambiae* during evolution.

Compared to the traditional taxonomic key method that relies on morphology for mosquito identification, our method is rapid due to its high throughput, making it ideal for mosquito surveillance programs. The best accuracy that we have reported here of 94% can be achieved and maintained by minimal training of the personnel involved. This performance is better than the morphology-based method, which has an average accuracy of 81% at the genus level but whose best and poorest performance can range from 100% to 50% depending on the expertise of the personnel (55). Furthermore, unlike the standard PCR assays, our method is rapid since it requires minimal sample preparation. It is also non-destructive and, after the initial costs of setting up the Raman microscope are taken into account, cost-effective because no chemical reagents are required. The Raman microscope used in this work costs about USD 100,000. However, it is a general-purpose system that is also used for other research projects in materials science, forensics, and bio-photonics. The beauty of Raman spectroscopy is that after the method development, a custom made, application-specific, hand-held system (56,57) can be designed with a preloaded library for mosquito identification. This will drastically decrease the initial cost of setting up a Raman system dedicated to mosquito identification to less than USD 30,000. The current cost of setting up a PCR system is about USD 40,000, with an expected constant requirement of reagents that may not be sustainable for laboratories in resource-limited settings. Our method also compares well with NIR spectroscopy (23,25,58,59), an optical technique with similar benefits to Raman spectroscopy. However, from a technical point of view, Raman measurements are made using laser light in (or close to) the visible range of the electromagnetic spectrum. Therefore, Raman spectroscopy is more appealing in the miniaturization of spectroscopy devices since visible light detectors are relatively cheaper than NIR detectors. Furthermore, Raman systems give better spatial resolution than NIR in spectral imaging applications (59). We believe that if Raman imaging is used, the classification accuracy achieved in this work may be improved.

## CONCLUSIONS

We have demonstrated the capability of Raman spectroscopy, in combination with machine learning algorithms, to discriminate medically important mosquito species: *Aedes aegypti, Anopheles gambiae* and *Culex quinquefasciatus*. The developed models have the potential to be extended to the discrimination of other insects.

The results suggest that a cuticular pigment, melanin, is responsible for discriminating the insect groups. A linear discrimination model, namely VT/PCA/LDA, performed moderately (85% accuracy; 69% sensitivity; 90% specificity) in discriminating the groups compared to VT/PCA/QDA, which exploited nonlinearity within the dataset, thus performing better (94% accuracy; 87% sensitivity; 96% specificity). This is the first time that a Raman spectroscopy method has been used to classify medically important mosquitoes.

Even though Raman spectroscopy gives complementary vibrational information to mid-IR spectroscopy and was, therefore, expected to detect signatures of cuticular lipids, the spectra were dominated by melanin spectral signatures. Melanins have traditionally been considered difficult to extract for chemical analysis due to their low solubility; hence they have not been explored in the classification of insects.

The classification models developed here were simple and limited to discrimination of mosquito species belonging to two medically important sub-families: Anophelinae and the Culicinae. They demonstrated the potential of Raman spectroscopy in insect classification. More complex classification models may need to be developed to classify morphologically indistinguishable species.

Raman spectroscopy coupled with an appropriate machine learning algorithm is, therefore, a potentially powerful tool for insect species discrimination and classification that could be used to identify morphologically indistinguishable cryptic species.

## Acknowledgement

We acknowledge the Swedish International Development Cooperation Agency (SIDA), through the International Science Programme (ISP), Uppsala University, for financial support.

